# Microbial species coexistence depends on the host environment

**DOI:** 10.1101/609271

**Authors:** Peter Deines, Katrin Hammerschmidt, Thomas CG Bosch

## Abstract

Organisms and their resident microbial communities form a complex and mostly stable ecosystem. It is known that the specific composition and abundance of certain bacterial species affect host health and fitness, but the processes that lead to these microbial patterns are unknown. We investigate this by deconstructing the simple microbiome of the freshwater polyp *Hydra*. We contrast the performance of its two main bacterial associates, *Curvibacter* and *Duganella*, on germ free hosts with two *in vitro* environments over time. We show that interactions within the microbiome but also the host environment lead to the observed species frequencies and abundances. More specifically, we find that both microbial species can only stably coexist in the host environment, whereas *Duganella* outcompetes *Curvibacter* in both *in vitro* environments irrespective of initial starting frequencies. While *Duganella* seems to benefit through secretions of *Curvibacter*, its competitive effect on *Curvibacter* depends upon direct contact. The competition might potentially be mitigated through the spatial structure of the two microbial species on the host, which would explain why both species stably coexist on the host. Interestingly, the fractions of both species on the host do not match the fractions reported previously nor the overall microbiome carrying capacity as reported in this study. Both observations indicate that the rare microbial community members might be relevant for achieving the native community composition and carrying capacity. Our study highlights that for dissecting microbial interactions the specific environmental conditions need to be replicated, a goal difficult to achieve with *in vitro* systems.

**Importance:** This work studies microbial interactions within the microbiome of the simple cnidarian, *Hydra*, and investigates whether microbial species coexistence and community stability depends on the host environment. We find that the outcome of the interaction between the two most dominant bacterial species in *Hydra*’s microbiome differs depending on the environment and only results in a stable coexistence in the host context. The interactive ecology between the host, the two most dominant microbes, but also the less abundant members of the microbiome, are critically important for achieving the native community composition. This indicates that the metaorganism environment needs to be taken into account when studying microbial interactions.

## Introduction

Eukaryotes form a distinct habitat for microbial communities (microbiomes) and these microbial associations are integral to life. The host with its associated microbial community, often dominated by bacteria but co-habited by fungi, protozoa, archaea, and viruses, is termed a metaorganism. Microbiomes can contain from few up to thousands of microbial species – the human microbiome, for example, is estimated to be comprised of about 5000 bacterial species (1-3). These host-associated microbial communities have been shown to enhance host function and contribute to host fitness and health (4). Changes in microbiome diversity, function, and density have been linked to a variety of disorders in many organisms (5-8).

A major goal in host-microbe ecology is to unravel the ecological and evolutionary dynamics of microorganisms within their communities. Of particular relevance are the factors that shape the stability and resilience of such communities, despite different fitness trajectories of the microbiome members. The microbial response to stress or perturbations, e.g. exposure to a new substrate, provides a selective advantage to certain members of the community. If the system cannot tolerate the change, the microbial community dramatically shifts until a different equilibrium state is reached (9). Frequency-dependent selection forces the host to adapt to these changes and select for or against the most frequent genotypes of their associated microbiota (10). There is, for example, strong evidence that species-specific antimicrobial peptides (AMPs) shape, control, and confine host-species specific bacterial associations (11, 12). In addition, microbial communities are not evenly distributed, e.g. along the gastrointestinal tract or between the lumen and the epithelial surfaces (2, 13, 14). These significant differences in niches or micro-habitats and their occupancy is known as spatial heterogeneity and will affect community assembly rules and dynamics (15, 16). Interspecies metabolic exchange is another key biotic force acting as a major driver of species co-occurrence in diverse microbial communities (17).

To experimentally address the composition and assembly of animal microbiomes current efforts have taken advantage not only of the traditional models such as the zebrafish, the fruit fly, and the nematode worm but also of other systems such as the honeybee, and crustacean species belonging to the genus *Daphnia* (18). All of these simple animal models can be raised and manipulated in the laboratory allowing for the discovery of fundamental principles of animal-microbiome interactions. As most of these models contain only a small number of taxa, a bottom-up approach can help to better understand these host-associated microbiomes using synthetic microbial communities (19, 20).

We here apply a reductionist approach to disentangle the inherent complexity of interactions in host-microbiomes. We use the freshwater polyp *Hydra vulgaris* and its microbiome, which has become a valuable experimental model in metaorganism research (21). *Hydra’s* ectodermal epithelial cells are covered with a multi-layered glycocalyx that provides a habitat for a species-specific and core microbiome of low complexity (11, 22, 23), from which most microbes can be cultured *in vitro* (23, 24). This allows the construction of synthetic communities of various complexities and contrasting the host (*in vivo)* to *in vitro* habitats (21). We focus on the two most abundant members of the microbiome that together constitute about 85% of *Hydra*’s simple microbiome, *Curvibacter* sp. AEP1.3 and *Duganella* sp. C1.2, (hereafter called *Curvibacter* and *Duganella*), where abundances of *Curvibacter* are several magnitudes higher as compared to *Duganella* (24). In this study, we want to understand the factors leading to this pattern of species coexistence and hypothesize that the respective environmental conditions are key for the outcome of microbial species interactions. After a characterization of the population dynamics of the two main colonizers when grown singly, we perform experiments where we combine host and non-host environments with invasion-from-rare set-ups to elucidate the effect of the host environment on microbial interactions. More specifically we test whether microbial species coexistence and community stability depends on the host environment.

## Results

### The *Hydra* ecosystem is characterized by an overall carrying capacity

Carrying capacity is defined as the maximum population size that an ecosystem can sustainably support without being degraded. This concept from macro-ecology can also be applied to host-microbe ecosystems. We here determine whether the *Hydra* ecosystem is characterized by a specific carrying capacity, and whether it can be reached again after the incubation of germ-free polyps with tissue homogenates of wild-type animals (conventionalized animals). We find that the carrying capacity of *Hydra* is highly stable among single *Hydra* wild-type polyps with 1.7* 10^5^CFUs per individual (standard deviation of ±0.3* 10^5^). This carrying capacity cannot be exceeded through the artificial addition of either *Curvibacter* or *Duganella* to wild-type polys. In the contrary, the addition of *Curvibacter*, leads to a significant reduction in overall microbial population size (Welch ANOVA, F_3_=7.054; P<0.005; Fig. 1A). Most importantly, the carrying capacity of wild-type and conventionalized polyps does not differ. These findings indicate the usability of germ-free polyps for the manipulation and construction of *in vivo* synthetic bacterial communities

**Fig. 1.**
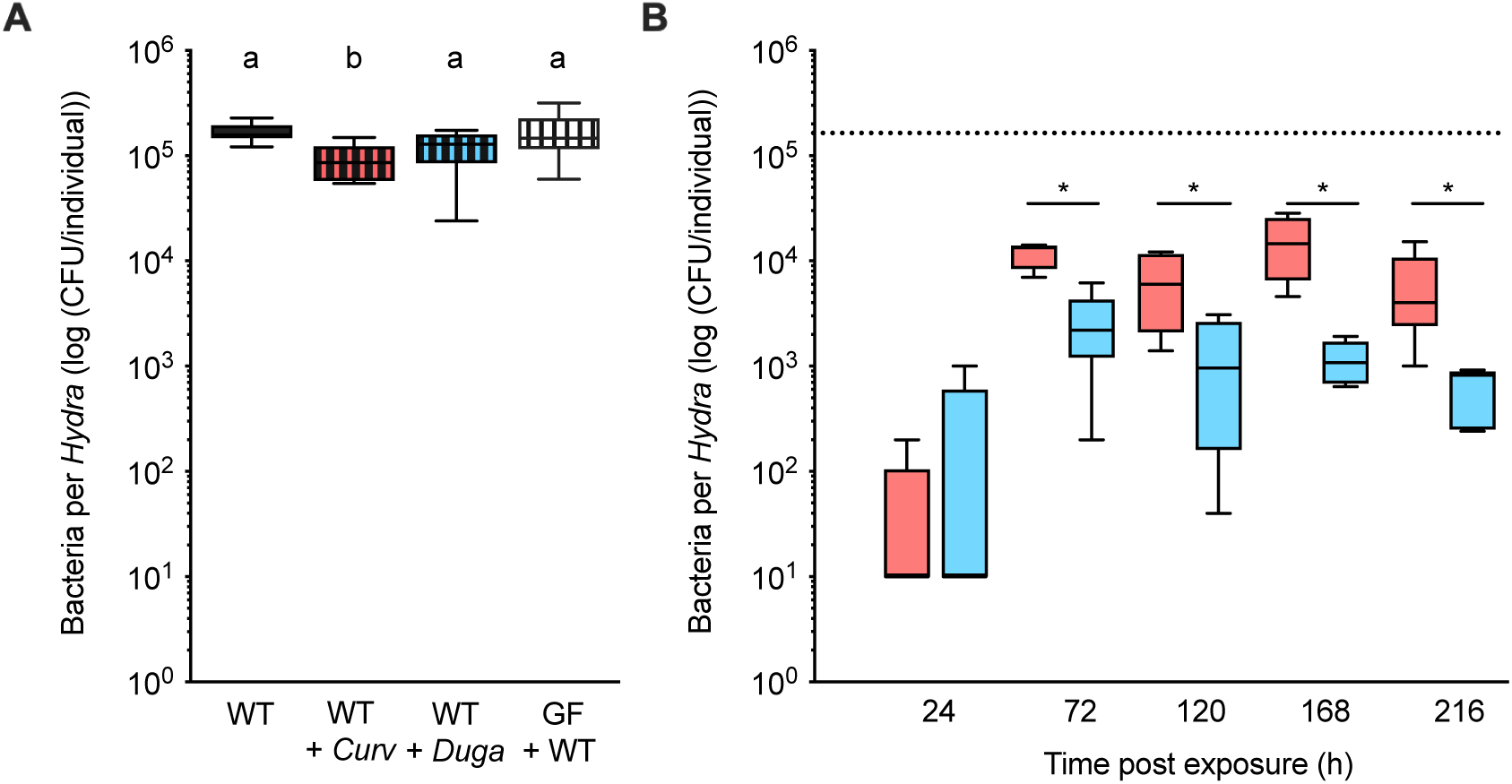
(A) Carrying capacity of the *Hydra* habitat in wild-type (WT) polyps, wild-type polyps and the addition of either the focal species *Curvibacter* (*Curv*) or *Duganella* (*Duga*) and germ-free (GF) animals incubated with native *Hydra* microbiota (conventionalised polyps) (each boxplot 16≥N≥6). (B) Time course analysis of microbial abundances in mono-associations of germ-free polyps with either *Curvibacter* (red) or *Duganella* (blue) (each boxplot N=6). The dashed line indicates the carrying capacity of wild-type polys.

### Species-specific carrying capacities for *Hydra’*s two main colonizers

In any habitat, the carrying capacity for each species is different and depends on several factors, including availability of nutrients, spatial distribution, and inter- and intra-species interactions. To assess the capabilities to colonize *Hydra*, we assess the population growth of *Curvibacter* and *Duganella* in mono-associations until they reach their individual carrying capacities. After about 72 h of growth on the host, both microbial species reach a stable population size (Fig. 1B). This carrying capacity, when grown singly on the host, differs between both strains, with *Curvibacter* reaching a higher population size than *Duganella* (estimated by post hoc contrasts; Generalized linear model: Full model: χ^2^=54.360, d.f.=9, P<0.0001; bacterial species × days post exposure: χ^2^=18.326, d.f.=4, P=0.0011). These significant differences last until the end of the experiment. The carrying capacity of mono-associations is about 10^4^ CFUs per individual for *Curvibacter*, whereas for *Duganella* the population size reaches on average only 1.5* 10^3^ CFUs per individual. Further, both mono-associations do not reach the overall carrying capacity of wild-type polyps. The variation in bacterial density between hosts is significantly higher in *Curvibacter* than in *Duganella* (Levene: F_1_=21.496, P<0.0001).

### Deconstructing the metaorganism: role of the host environment and of the most dominant co-colonizing microbial species

Here we perform an experiment to deconstruct some of the interactions within the *Hydra* metaorganism by (i) exploring the role of the host as a microbial habitat through the comparison of microbial population dynamics *in vivo* with *in vitro* environments, and (ii) determining the role of the second most-dominant co-colonizer within the *Hydra* microbiome through performing di-association experiments with different starting frequencies of both microbial species. The previous experiment showed that the carrying capacity for each species was reached after 72 h post inoculation and stably maintained thereafter. Here we adjust our sampling intervals accordingly: we focus on the critical period that determines the outcome of the colonization process, i.e. the first 96 h post inoculation, and shorten the intervals in-between sampling to 12 h. This should allow for a detailed monitoring of the microbial population dynamics until the respective carrying capacities are reached.

#### The effect of the environment on microbial growth kinetics in mono-associations

To determine the relative importance of the host for microbial population dynamics and community stability, we chose two *in vitro* environments to contrast to the host: static, which closely resembles the host habitat in that it provides spatial heterogeneity facilitating bacterial interactions and mixed, where direct interactions between individual bacteria cannot be established but where individual bacteria have (unlimited) access to resources and oxygen.

We find growth rates of *Curvibacter* not to significantly differ between the host and the microcosm environments. This is in marked contrast to *Duganella*, where significantly higher growth rates were observed in the non-host as compared to the host environment. In all environments, except for the host, *Duganella* achieved a significantly higher growth rate than *Curvibacter* (determined by post hoc t-tests; ANOVA: *R*^2^=0.827; Full model: *F*_5,15_ = 14.333; P<0.0001; bacterial species × environment: *F*_2_ = 15.592; P=0.0002; Fig. 2).

**Fig. 2.**
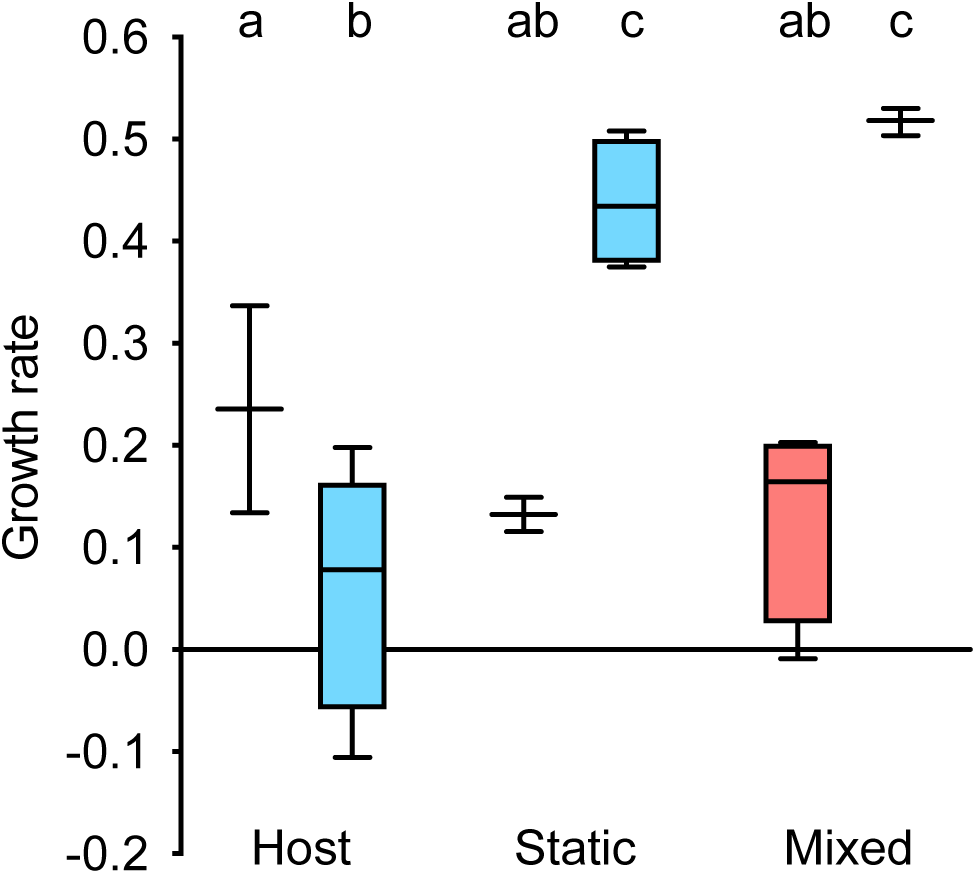
Bacterial growth rates of *Curvibacter* (red; each boxplot 4≥N≥2) and *Duganella* (blue; each boxplot 6≥N≥3) in mono-associations are habitat dependent. Compared are the host habitat (*in vivo*), and two *in vitro* environments: heterogeneous (static microcosms), and homogenous (mixed microcosms).

#### The effect of microbial interactions on microbial performance varies depending on the environment

Here we test whether microbial performance is affected by the presence of the other, most dominant co-colonizer from the *Hydra* microbiome. Again, we contrast the *in vivo* environment with the two *in vitro* environments to be able to test whether the environment, the fellow microbes or an interaction of both determines microbial population dynamics and community stability.

The overall population dynamics in di-associations resembles the one observed in mono-associations, namely that irrespective of the environment, the carrying capacity in all habitats is reached at about 72 h after inoculation. Both microcosm environments are characterized by a carrying capacity of 10^7^-10^8^ CFUs/ml, and so exceed the *in vivo* carrying capacity by a factor of 10^4^ (Fig. 3). Nevertheless, di-associations on the host also fail to reach the overall carrying capacity of wild-type polyps (Welch ANOVA, F_1_=441.929; P<0.001) and reach a comparable carrying capacity as in the mono-colonisations of *Curvibacter* and *Duganella* (ANOVA: *F*_2,44_ = 2.011, P=0.146). Both bacterial species do not match the species-specific carrying capacities as measured in mono-colonisations on the host: whereas *Curvibacter* fails by a power of 10 to reach its density in the mono-colonisations, *Duganella* outgrows it by a power of 10.

**Fig. 3.**
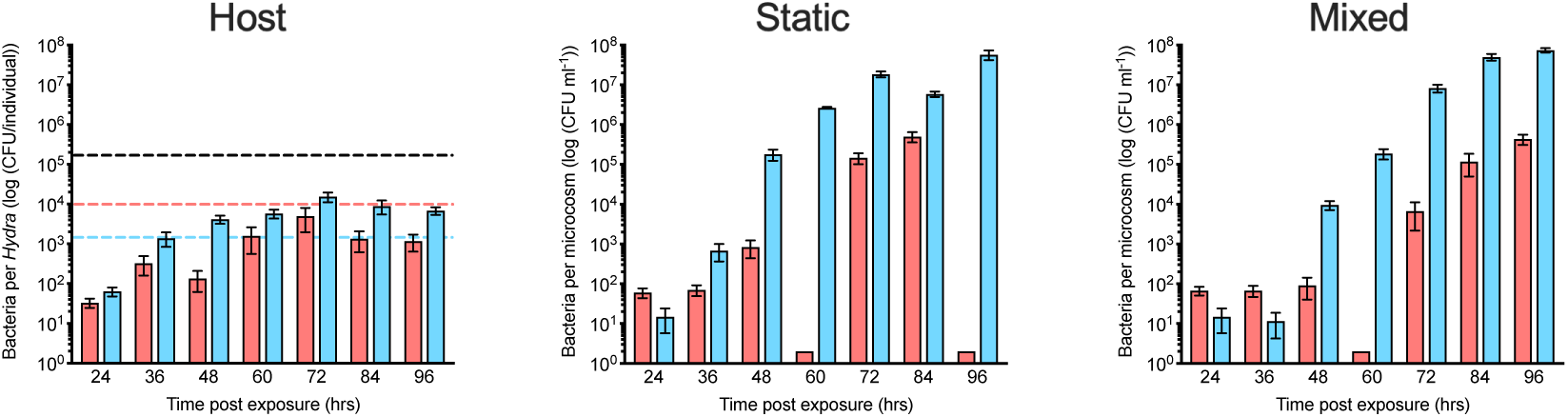
Carrying capacity of the *in vivo* and *in vitro* habitats used in this study. Shown are pooled total numbers of colony forming units (CFUs) from all di-association experiments with *Curvibacter* (red) and *Duganella* (blue) (shown are s.e.m. based on 18≥N≥11 for the host, 12≥N≥9 for static, and 12≥N≥4 for mixed). The dashed black line indicates the carrying capacity of WT polyps, the dashed red line the species-specific carrying capacity of polyps during *Curvibacter* mono-associations and the dashed blue line the species-specific carrying capacity of polyps during *Duganella* mono-associations.

To control for frequency-dependent microbial population dynamics, we competed *Curvibacter* and *Duganella* in three different starting frequencies (Fig. 4). In both non-host environments, *Duganella* outcompetes *Curvibacter* within 48 h post exposure. From then onwards, frequencies of *Curvibacter* are low, reaching a maximum of about 10%. This pattern does not depend on the initial frequency at the start of the experiment. Most interestingly, this pattern is not observed in the host environment: here, a decrease in the fraction of *Curvibacter* can be observed in all three initial frequencies but never to a point where it cannot be detected in the population. From 72 h post exposure onwards the population on the host has reached a stable state, with *Curvibacter* making up 20% of the total bacterial population.

**Fig. 4.**
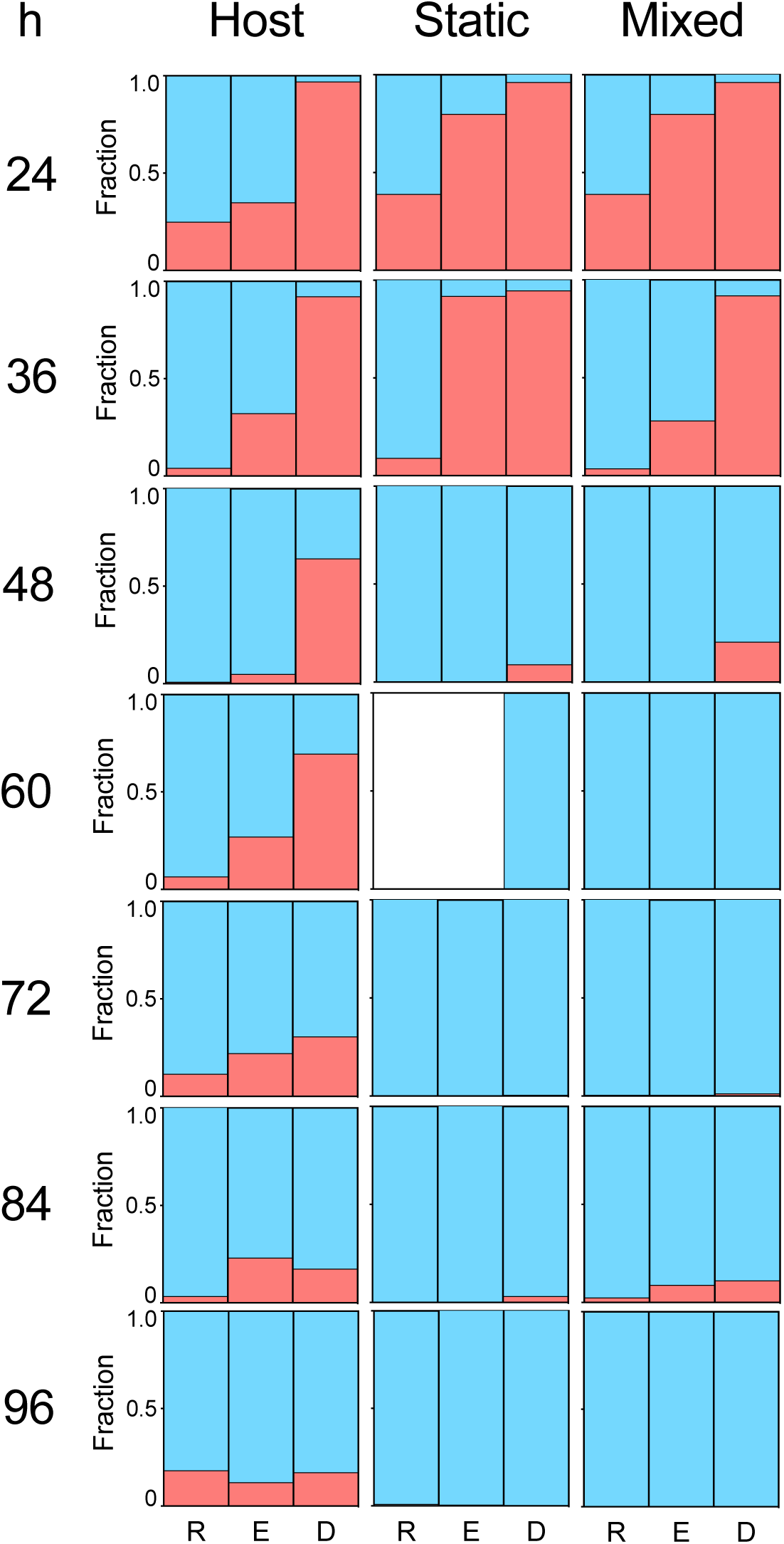
Time course of fractions of *Curvibacter* (red) and *Duganella* (blue) in the three different habitats obtained from di-association experiments. The initial inoculation frequency of *Curvibacter* varied from being rare (R), to equal (E), to dominant (D) in comparison to *Duganella* (each bar 6≥N≥3; except for: Host, D, 24 h, where N=2, and for Static, R and E 60 h, where N=0 due to contamination of plates).

### Zooming in on the interaction between the two most dominant microbes in the *Hydra* microbiome

In every environment, *Duganella* had a negative impact on *Curvibacter*, while the presence of *Curvibacter* led to an increased *Duganella* carrying capacity in both, host and microcosm environments. We here performed growth assays of both microbial species in spent medium (cell-free supernatant) of the other microbial species to test whether contact dependent or contact independent interactions determined the observed population dynamics. Interestingly, we did not observe reduced growth of *Curvibacter* in the supernatant of *Duganella*, but the opposite, with *Curvibacter* growing to higher abundances in cell-free supernatant as compared to abundances in *Hydra* medium (ANOVA: *R*^2^=0.924, *F*_1,8_ = 97.312, P<0.0001; Fig. 5A). Also, *Curvibacter* supernatant led to higher *Duganella* abundances as compared to *Hydra* medium (ANOVA: *R*^2^=0.946, *F*_1,8_ = 140.628, P<0.0001) – but only after a significantly longer lag time (ANOVA: *R*^2^=0.996, *F*_1,8_ = 1876.566, P<0.0001; Fig. 5B).

**Fig. 5.**
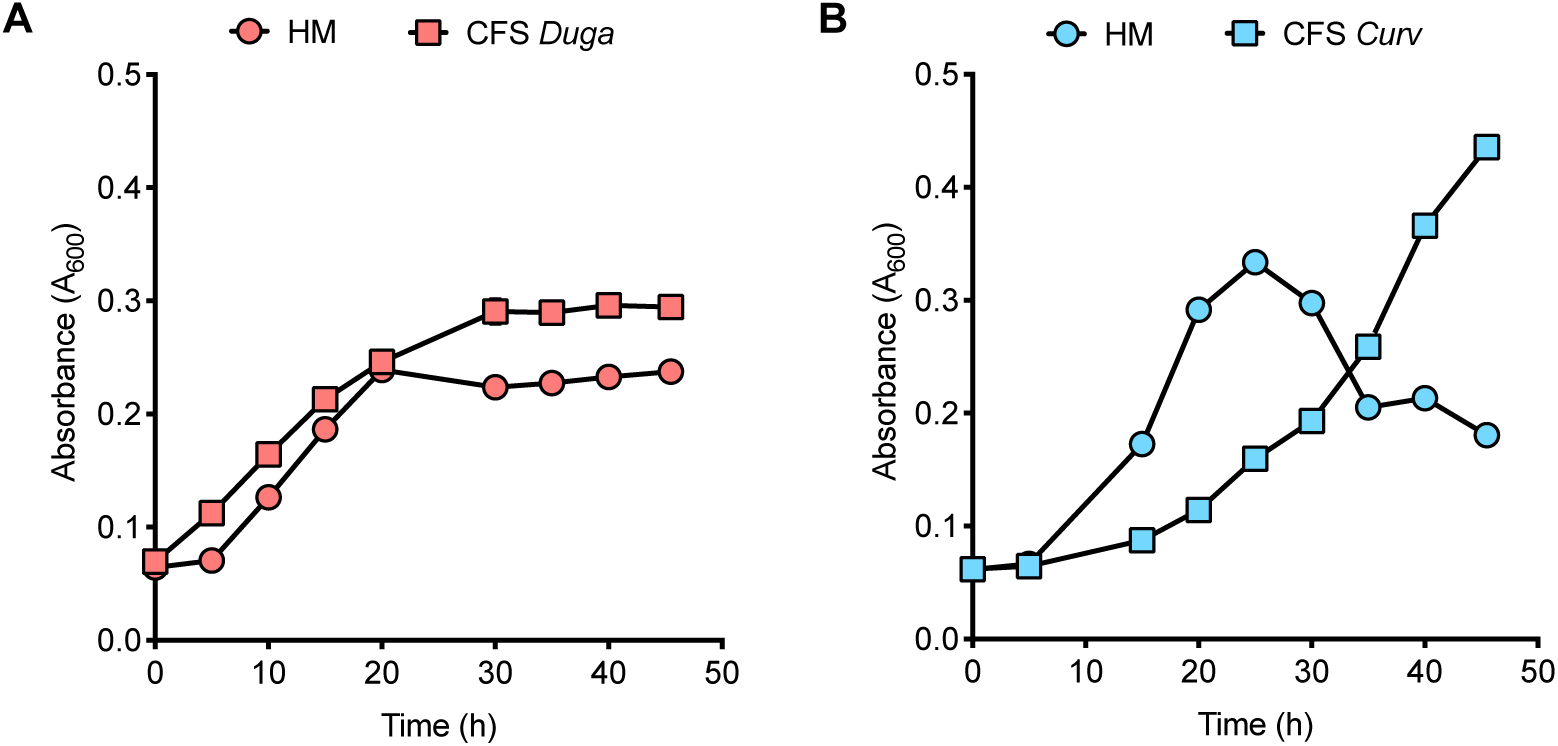
Bacterial growth dynamics of *Curvibacter* (red; A) and *Duganella* (blue; B) in *Hydra* medium (HM) and in the presence of *Duganella* (CFS *Duga*) or *Curvibacter* (CFS *Curv*) cell-free supernatant (CFS). Plotted are means (± s.e.m. based on N=5).

## Discussion

One of the major challenges in microbiome research is to understand the factors that influence the dynamics and stability of host-associated microbial communities. Of particular relevance for this are the processes governing assembly (25, 26) and resilience (27). Insights into such processes in bacterial populations within their native host environments can be gained through a number of ways. A bottom-up approach, for example, allows to elucidate the basic principles of community assembly as has been shown in non-host associated microbial communities (28, 29). This strategy is becoming increasingly popular in metaorganism research and has produced informative results, such as in the zebrafish (30) or the nematode *Caenorhabditis elegans* (19). Here we use *Hydra* and its microbiome for ‘deconstructing’ a metaorganism and its interactions (21). To determine the relative importance of the host on the interactions of *Curvibacter* and *Duganella*, we performed mono- and di-association experiments *in vivo* and in two i*n vitro* environments (Fig. 6). As community structure can be influenced by initial species abundances (31), we also performed all di-association experiments using various initial starting frequencies. Interestingly we found that in the *Hydra* habitat *Curvibacter*, independent of its inoculation frequency, and after the initial establishment period of 72 h, reached a constant fraction of about 20%, whereas it was only present at very low frequencies or went extinct in both *in vitro* habitats (see also (32) for similar patterns in a homogeneous *in vitro* environment). Wright and Vetsigian (33) recently demonstrated in pairwise competitions between bacteria of the genus *Streptomyces* that the winner is often the species that starts at high initial abundance. We find that ‘survival of the common’ does not apply to *Curvibacter* in a non-host environment, whereas pairwise competitions in the host habitat show signs of stabilisation between *Curvibacter* and *Duganella*. This suggests that within the host environment both strains can stably coexist, which is in contrast to the *in vitro* environment, where we find competitive exclusion (*Duganella* excludes *Curvibacter*). Nevertheless, the resulting fractions in the *in vivo* di-association experiments (*Curvibacter* 20% and *Duganella* 80%) do not represent the species fractions found in wild-type polyps. Here *Curvibacter* represents 75% and *Duganella* 11% of the whole community (24), which indicates that the rare microbiome members might be relevant for achieving the native community composition. When *Curvibacter* and *Duganella* are introduced separately to the host (in mono-associations), each bacterial species is capable to robustly colonize the host to high abundances. This confirms earlier findings from Wein *et al*. (34) that *Curvibacter* is able to reach stable abundances on the host. A similar observation has been made for *Aeromonas* and *Vibrio* colonizing patterns of the gut of larval zebrafish (35) and of microbes colonizing the gut of *C. elegans* (19). While in mono-associations *Curvibacter* reaches higher abundances than *Duganella*, the opposite is true for the di-associations. Here, *Curvibacter* fails to reach its species-specific carrying capacities by a factor of 10 (as compared to the mono-association), *Duganella* outgrows it by a factor of 10.

**Fig. 6.**
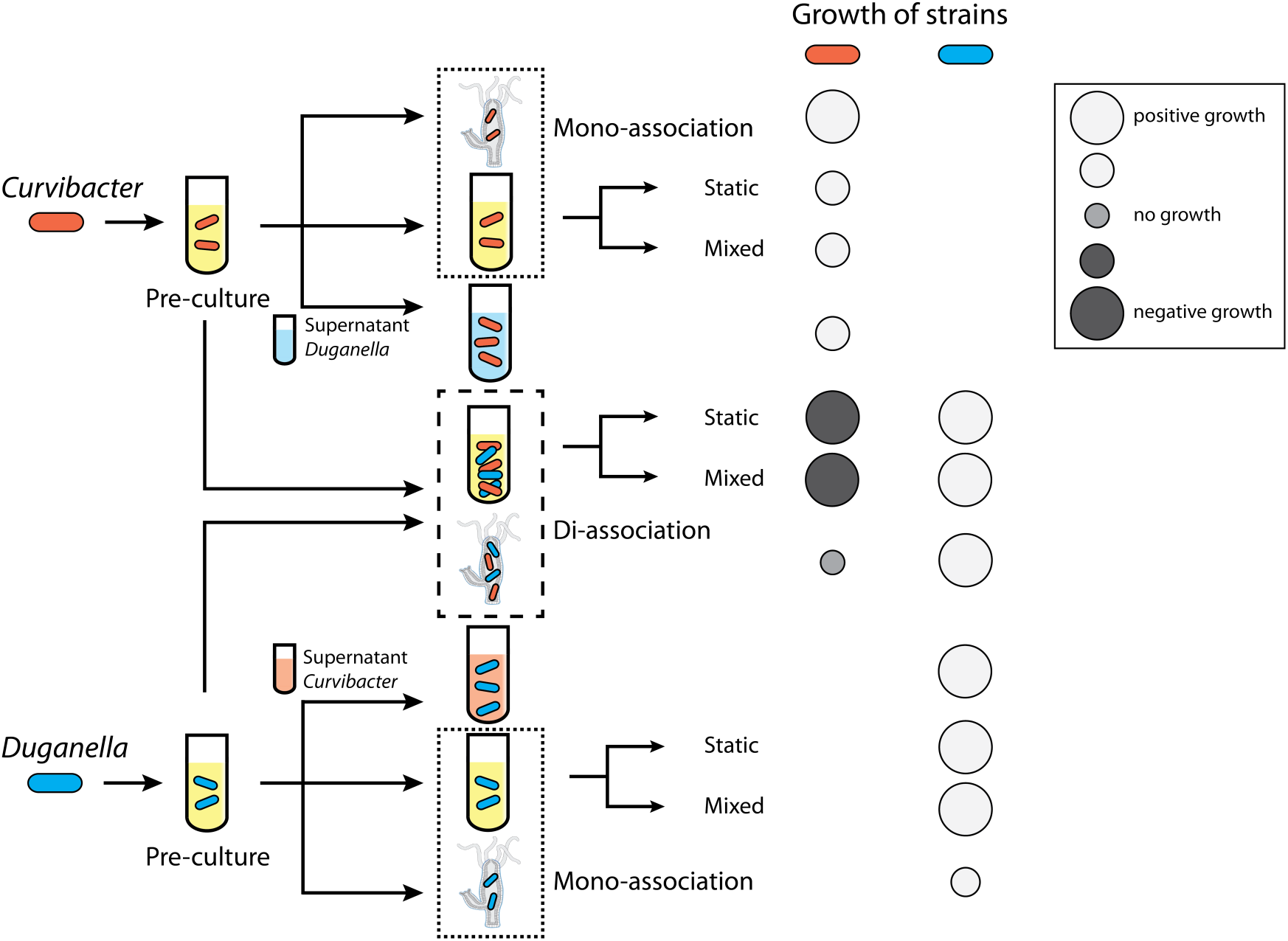
Overview of performed experiments and effects on growth of *Curvibacter* (red) and *Duganella* (blue).

Results from the di-association experiments clearly show that the abundances and relative frequencies of *Curvibacter* and *Duganella* as measured in wild-type *Hydra* cannot be explained by their interactions in the host context alone as this led to a frequency reversal, making *Duganella* more abundant than *Curvibacter. Duganella* however reaches comparable carrying capacities as measured in wild-type *Hydra* polyps. These observations indicate that the less frequent community members (each 2% and less) are important for achieving the overall carrying capacity of the *Hydra* microbiome. We hypothesise that two aspects are of importance here – (i) the low abundant microbes might be able to utilize different resources as compared to *Curvibacter* and *Duganella* and so inhabit different ecological niches within the microbiome, which the two main colonizers cannot fill and (ii) they likely interact in a positive way (either directly or indirectly) with (at least) *Curvibacter*, enabling it to reach higher carrying capacities. Evidence for the importance of rare-species comes from the human intestinal microbiota, which contains many low-abundance species (36) with some of them having a large impact on inducing dysbiosis in the microbiome and on guaranteeing host health (37, 38). It is thus important to note that *Hydra*’s carrying capacity is not solely determined by the host (resources) alone but also by the interactions within the microbiome. In microbiome research, the significance of an overall host carrying capacity has been largely overlooked until very recently where a link between host health and microbiome density has been reported (8). Bacterial levels have been quantified for only a few model organisms such as in the gut of larval zebrafish (39) or the gut of *Drosophila melanogaster* (40). We here show that also wild-type *Hydra* is characterized by an overall carrying capacity of about 10^5^ bacteria per polyp that is stable in adult polyps and can be artificially assembled through the re-population of germ-free animals with the native microbial community. This is an important prerequisite for conducting the *in vivo* experiments, where colonization patterns of single species from *Hydra’s* microbiome are individually followed.

To investigate the interaction between *Curvibacter* and *Duganella* in more detail, we tested whether the interactions between the two species are contact dependent by performing spent medium assays. Our results indicate that the effect of *Duganella* on *Curvibacter* might depend on direct contact, as *Duganella* supernatant did not negatively affect the growth of *Curvibacter*. In contrast, *Curvibacter* supernatant led to an initial time lag in *Duganella* growth, which was followed by an exponential growth phase after about 35 h (note that such a time lag is not visible in the other media). The same pattern can be observed in the fractions in the di-association experiments – also here, *Duganella* started to outcompete *Curvibacter* only after an initial delay of 36 h (Fig. 4). It is interesting to speculate what might lead to this pattern. The fact that this also becomes apparent in the supernatant experiments suggests that it is mediated by products in the supernatant of *Curvibacter*, which might metabolically be not directly assessable, so *Duganella* needs to adjust its physiology accordingly.

The observation that *Curvibacter* is not negatively affected by the *Duganella*’s supernatant suggests that direct contact is needed between the two species for *Duganella* to outcompete *Curvibacter*. The competition can be either passive, where strains compete for the same resources or active, where strains directly harm one another (41). Thus, one explanation for the stable coexistence of the two species on *Hydra* could be that the host environment leads to a (partial) spatial segregation of *Hydra*’s most dominant colonizers, reducing between-species contact, as has been shown for biofilms (42). For *Hydra* it is known that it shapes its microbiota through the secretion of antimicrobial peptides (43) and neuropeptides (44), which influences the microbial spatial structure. In addition, other host mechanisms have been predicted, such as the provisioning of carbon sources via epithelial feeding or releasing of specific adhesive molecules from epithelial surfaces targeted at specific microbes (45). Thus, it is important to conclude that while stability of microbial communities depend on interactions between different bacterial strains and species, these interactions need to occur in their native environment, the host.

## Materials and Methods

### Animals used, culture conditions and generation of germ-free animals

*Hydra vulgaris* (strain AEP) was used for carrying out experiments and cultured according to standard procedures at 18°C in standardized culture medium (*Hydra* medium (HM)) (46). Animals were fed three times a week with 1st instar larvae of *Artemia salina*. Germ-free (GF) polyps were obtained by treating wild-type (WT) animals with an antibiotic cocktail solution containing 50 *μ*g/ml ampicillin, neomycin, streptomycin, rifampicin and 60 *μ*g/ml spectinomycin as previously described (43, 47). The antibiotic cocktail solution was exchanged every 48 h and the antibiotic treatment lasted for two weeks, after which polyps were transferred into antibiotic-free sterile HM for recovery (four days). The germ-free status of polyps was confirmed as previously described (43). During antibiotic treatment and re-colonization experiments, polyps were not fed.

### Bacterial strains and media

The bacterial strains used in this study are *Curvibacter* sp. AEP1.3 and *Duganella* sp. C1.2., which have been isolated from *Hydra vulgaris* (strain AEP) (24). These bacteria were cultured from existing isolate stocks in R2A medium at 18°C, shaken at 250 r.p.m for 72 h before use in the different experiments. R2A was chosen as medium as it was used to isolate bacterial strains and to be able to compare results to previous published *Hydra* papers (24, 32).

### Carrying capacity of the host

To determine the carrying capacity of the *Hydra* habitat the microbial load of individual *Hydra* polyps (N=16) was determined. In addition to wild-type polyps the carrying capacity of conventionalized polys (N=12), obtained by incubating germ-free polyps with tissue homogenates of wild-type animals (per germ-free polyp one wild-type polyp was used) for 24 h was also determined. To test whether the carrying capacity can artificially be increased or destabilises upon self-challenge we added either *Curvibacter* or *Duganella* to wild-type polyps (N=6) (approximately 5×10^3^ cells for 24 h). After incubation all polyps were washed with and transferred to sterile HM and further incubated at 18°C and sampled after 120 h. Polyps were first washed three times with sterile HM to remove non-associated bacteria and then transferred to an Eppendorf tube containing sterile HM. After homogenization using a sterile pestle serial dilutions of the homogenate were plated on R2A agar plates to determine colony-forming units (CFUs) per individual.

### Tracking microbial mono-associations in *Hydra* over time

Germ-free polys were inoculated in their aquatic environment with single bacterial strains (mono-associations). Individual germ-free polyps were incubated with 5×10^3^ cells of *Curvibacter* or *Duganella* in 1.5 ml Eppendorf tubes containing 1 ml of sterile HM. After 24 h of incubation all polyps were washed with and transferred to sterile HM, incubated at 18°C and followed over a period of 216 h. For each treatment 6 polys per time point were independently analysed. Every 48 h individual polyps were collected to determine CFUs a described above.

### Microbial growth kinetics of mono- and di-associations *in vivo* and *in vitro*

To study the initial phase of colonization, i.e. 96 h post inoculation (see Fig. 1B) in more detail microbial growth of *Curvibacter* and *Duganella* was determined in different habitats; the host habitat (*in vivo*) and two different microcosm environments (*in vitro*). The static incubation provided a spatially structured habitat (heterogeneous), whereas shaking of the microcosms (mixed treatment) eliminated the spatial structure (homogenous).

#### Mono-associations

All germ-free polyps and microcosms were inoculated from the same bacterial inoculation culture with approximately 5×10^3^ cells of *Curvibacter* or *Duganella* for 24 h, washed with and transferred to sterile HM. Samples were taken every 12 h for 96 h. For *Hydra* six polyps were sacrificed at each time point and colony-forming units (CFUs) were determined as described above. As microcosms 24-well plates were used. Wells were filled with 2 ml of R2A medium, inoculated and incubated at 18°C either under static or shaken (200 r.p.m) conditions. Each time point was replicated four times and serial dilutions were plated on R2A agar plates to determine CFUs. Growth rates of each strain (A and B) were determined for the exponential growth phase (36-48 h) and were calculated as g = ln (A_f_/A_0_) and g = ln (B_f_/B_0_), where A_0_, B_0_ is the starting density at time 0 and A_f_, B_f_ is the final density at time *f*.

#### Di-associations

Density dependent competiveness fitness assays of the two most dominant colonizers *Curvibacter* and *Duganella* were performed *in vivo* and *in vitro*. The same host and microcosm experiments as described above were performed except for using microbial di-associations of *Curvibacter* and *Duganella* with the frequency of *Curvibacter* being rare (10:90), equal (50:50), or dominant (90:10). As *Curvibacter* and *Duganella* form distinct colonies on R2A agar plates, their frequency can be determined by plating serial dilutions (32). Six polyps and four microcosm replicates were assayed per treatment (static and mixed) and time point. Also this data allowed determining the different carrying capacities of the *in vivo* and *in vitro* habitats used. Growth rates of each strain were calculated as above (Fig. S1).

### Spent medium assay to measure interaction activity between microbial strains

Spent medium (cell-free supernatant (CFS)) of both microbial strains *Curvibacter* and *Duganella* was prepared by growing them for 72 h in R2A medium at 18°C (shaken at 250 r.p.m) until stationary phase was reached. Cultures were then centrifuged at 1000 g for 20 min and the supernatant was passed through a 0.22 µm filter. Growth assays consisted of mixing 100 µl of a culture of *Curvibacter* or *Duganella* (adjusted to an OD_600_ of 0.025) with the corresponding supernatant (100 µl) of the other strain. As control the same volume of HM was added to the strains instead of the supernatant. Growth kinetics were examined in a 96-well plate using a Tecan Spark 10M microplate reader. The plate was incubated at 18°C and moderately shaken for 10 sec prior to each read. Absorbance at 600 nm was measured every 30 min over a period of 48 h, and data from 5 h intervals are plotted for clarity. Each treatment was replicated 5 times.

### Statistical analysis

A Welch ANOVA (and subsequent Dunnett’s posthoc test) was used to test for differences in bacterial abundance patterns (‘bacteria per *Hydra*’) in wild-type versus manipulated hosts as variances between the different groups were not equally distributed.

Differences during mono-colonizations of *Curvibacter* and *Duganella* over time were assessed using a Generalized linear model (error structure: normal; link function: identity). The response variable was ‘bacteria per *Hydra*’, and explanatory variables were ‘bacterial species’, ‘time’ and ‘bacterial species’ × ‘time’. Differences between the two bacterial species on each day were detected with post hoc contrasts.

Carrying capacities were compared between wild-type *Hydra* and di-associations using a Welch ANOVA, and between mono- and di-associations using an ANOVA.

Analysis of variance (ANOVA) and subsequent post hoc t-tests were used to test for differences in growth rates of the two competitors when grown singly in the different environments. The response variable was ‘growth rate’, and explanatory variables were ‘bacterial species’, ‘environment’ and ‘bacterial species’ × ‘environment’.

Differences in the growth rates in the di-associations of *Curvibacter* and *Duganella* in the different environments and dependence on initial frequency were assessed using a Generalized linear model (error structure: normal; link function: identity) and post hoc contrasts. For each bacterial species, a separate model was calculated with the response variables being either ‘growth rate *Curvibacter*’ or ‘growth rate *Duganella*’, and the explanatory variables were ‘environment’, ‘starting density’ and ‘environment’ × ‘starting density’.

Differences between the maximal bacterial abundances of both species and lag time of *Duganella* when grown in the supernatant of the respective other species as compared to growth in *Hydra* medium were assessed using ANOVA.

Sample size was chosen to maximise statistical power and ensure sufficient replication. Assumptions of the tests, that is, normality and equal distribution of variances, were visually evaluated. Non-significant interactions were removed from the models. All tests were two-tailed. Effects were considered significant at the level of P < 0.05. All statistical analyses were performed with JMP 9. Graphs were produced with GraphPad Prism 5.0, and Adobe Illustrator CS5.1.

## Acknowledgements

PD received funding from the European Union’s Framework Programme for Research and Innovation Horizon 2020 (2014–2020) under the Marie Skłodowska-Curie Grant Agreement No. 655914 and KH under the Marie Skłodowska-Curie Grant Agreement No. 657096. Both also received a Reintegration Grant from the Deutscher Akademischer Austausch Dienst (DAAD). This work was further supported by the Deutsche Forschungsgemeinschaft (DFG) Collaborative Research Center (CRC) 1182 (“Origin and Function of Metaorganisms”). TB gratefully appreciates support from the Canadian Institute for Advanced Research (CIFAR) and thanks the Wissenschaftskolleg (Institute of Advanced Studies) in Berlin for a sabbatical leave. We thank Chaitanya Gokhale for feedback on the manuscript.

---

**Fig. S1.**
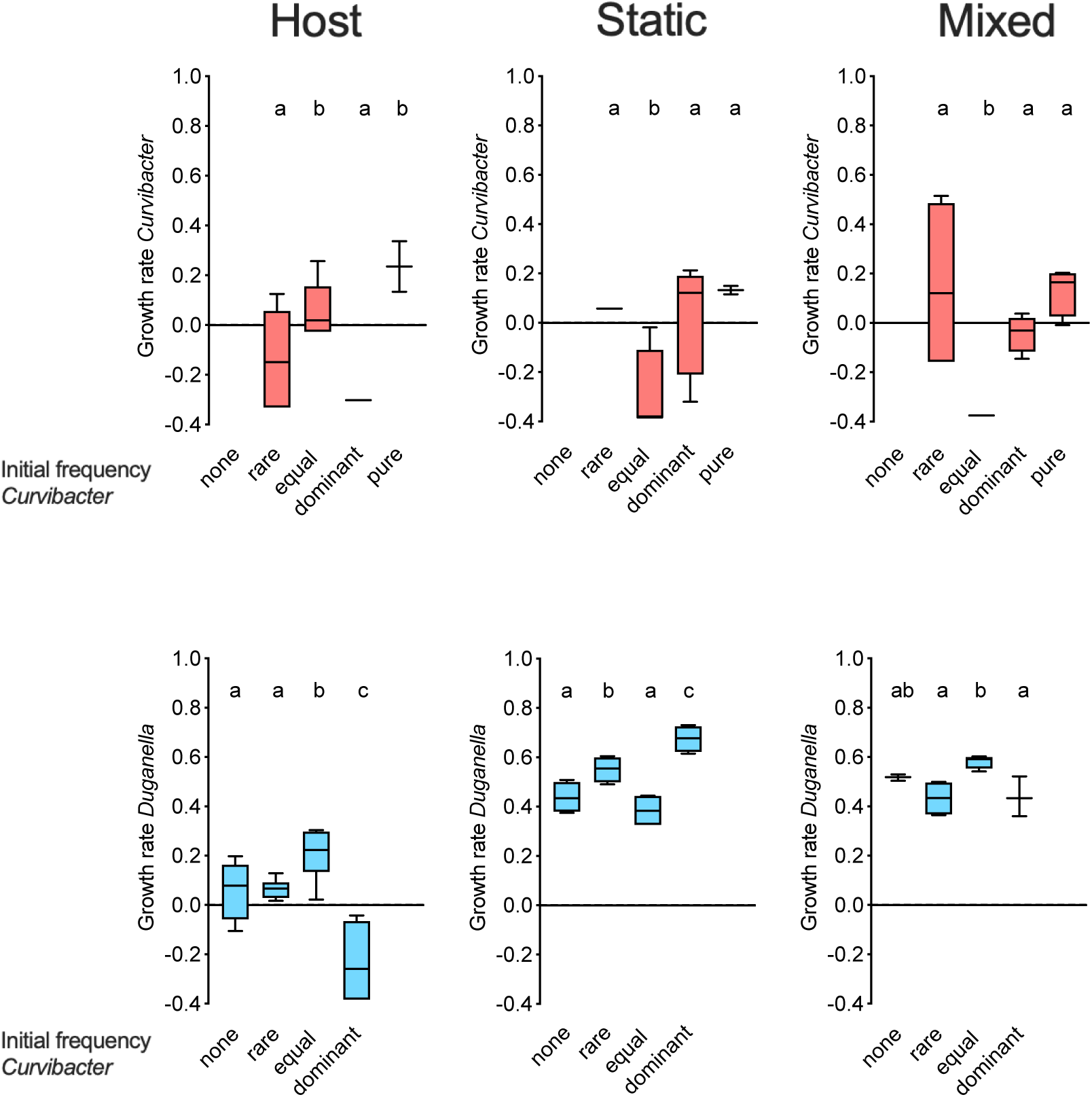
Bacterial growth rates of *Curvibacter* (red) and *Duganella* (blue) from mono- and di-association experiments across the different habitats and initial frequencies tested (each boxplot 4≥N≥3). Overall *Curvibacter* growth rate in di-associations are lower or not different from the mono-associations (as estimated by post hoc contrasts; Generalized linear model: Full model: χ2=45.790, d.f.=11, P<0.0001; environment × initial frequency: χ2=33.685, d.f.=6, P<0.0001). *Curvibacter* grows significantly differently when inoculated in equal densities as compared to the rare and dominant starting frequencies across the different environments. Whereas in the host, *Curvibacter* grows better when in equal density with *Duganella*, the opposite is true for both *in vitro* environments. As observed for the growth of *Duganella* in mono-colonisations, growth rates are always higher in the non-host environments irrespective of initial frequency (Generalized linear model: Full model: χ2=130.278, d.f.=11, P<0.0001; environment × initial frequency: χ2=59.723, d.f.=6, P<0.0001). Whereas, in di-associations, negative growth rates can be detected only once for *Duganella*, it happens more frequently in the *Curvibacter*, indicating a direct or indirect negative effect of *Duganella*.

